# A cell-based scrambling assay reveals phospholipid headgroup preference of TMEM16F on the plasma membrane

**DOI:** 10.1101/2025.06.25.661602

**Authors:** Chin Fen Teo, Sami T. Tuomivaara, Niek van Hilten, David Crottès, Yuh Nung Jan, Michael Grabe, Lily Y. Jan

**Affiliations:** Howard Hughes Medical Institute, University of California, San Francisco, CA 94143, USA; Department of Physiology, University of California, San Francisco, CA 94143, USA; Department of Obstetrics, Gynecology, and Reproductive Sciences, Center for Reproductive Sciences, Eli and Edythe Broad Center for Regeneration Medicine and Stem Cell Research, Sandler-Moore Mass Spectrometry Core Facility, University of California, San Francisco, CA 94143, USA; Department of Pharmaceutical Chemistry, Cardiovascular Research Institute, University of California, San Francisco, CA 94143, USA

**Author notes:** Meilahti Proteomics Unit, Department of Biochemistry and Developmental Biology, Faculty of Medicine, Helsinki Institute of Life Science, University of Helsinki, 00014 Helsinki, Finland (Sami T. Tuomivaara). Inserm UMR 1069 N2Cox, Niche, Nutrition, Cancer & Métabolisme Oxydatif, Tours, France. **Corresponding Author:** Lily Y. Jan **Email:**.

**Keywords:** TMEM16F, phospholipid scrambling, fluorescence polarization, plasma membrane, coarse-grained molecular simulations

## Abstract

The asymmetric resting distribution of the three major phospholipid classes on the mammalian plasma membrane, with phosphatidylserine and phosphatidylethanolamine mostly on the inner leaflet, and phosphatidylcholine mostly on the outer leaflet, is maintained by ATP-dependent flippases and floppases that exhibit headgroup selectivity. Upon signaling cues, this asymmetry can be dissipated by various phospholipid scramblases, allowing cells to respond to stimuli and adapt to different physiological contexts. The prevailing view in the field is that phospholipid scramblases on the plasma membrane act without headgroup preference. Here we report contrary experimental evidence based on a phospholipid scrambling assay that quantifies the fluorescence polarization of nitrobenzoxadiazole-labeled phospholipids for kinetic monitoring of phospholipid scrambling on the plasma membrane of living cells. Our experiments reveal that the plasma membrane-residing calcium-activated phospholipid scramblase TMEM16F preferentially acts on phosphatidylserine and phosphatidylcholine over phosphatidylethanolamine.

**Significance Statement:** Phospholipid scramblases on the mammalian plasma membrane are thought to act promiscuously without preference for headgroup. Thoroughly addressing this question, however, requires the development of new methodologies. We devised a cell-based phospholipid scrambling assay that utilizes the fluorescence polarization of nitrobenzoxadiazole (NBD)-labeled phospholipids, allowing the monitoring of their scrambling in a native environment. We discovered that the plasma membrane-residing calcium-activated phospholipid scramblase TMEM16F preferentially acts on phosphatidylserine and phosphatidylcholine over phosphatidylethanolamine.

**Major Classification**: Biological Sciences; **Minor Classification:** Cell biology, Biophysics and Computational Biology

## Introduction

The asymmetric distribution of phospholipids (PLs) on the plasma membrane is collaboratively maintained by several classes of ATP-dependent flippases and floppases that exhibit PL headgroup selectivity (1). The resting distribution of three major PL classes on the plasma membrane, with phosphatidylserine (PS) and phosphatidylethanolamine (PE) mostly on the inner leaflet and phosphatidylcholine (PC) mostly on the outer leaflet, is critical in sustaining mammalian cellular physiology (2–4). Of equal importance is the regulated reduction of this asymmetry by different classes of ATP-independent PL scramblases (4–9), including those in the apoptosis-activated Xkr family (10, 11), the calcium-responsive TMEM16 family (12, 13), as well as the mechanosensitive TMC (14, 15) and TMEM63 families (16). These PL scramblases mediate crucial physiological events such as exposing an “eat-me” signal for apoptotic clearance or fine-tuning plasma membrane fluidity, curvature, and tension to enable various plasma membrane processes (2–5, 7–9). Unlike flippases and floppases, which utilize ATP to maintain the asymmetric distribution of PLs in the lipid bilayer, scramblases harness the potential energy stored in the asymmetry itself, driving towards PL equalization on the two leaflets along the concentration gradient (2, 3, 8).

Ubiquitously expressed and plasma membrane-localized Transmembrane Protein Family 16 Member F (TMEM16F), also known as anoctamin-6 (ANO6), is one of the best-characterized calcium-activated PL scramblases (4–6, 8, 9). Originally identified as a key determinant in the PS exposure and extracellular vesicle release from platelets for blood coagulation (13) and the genetic contributor to the rare hereditary bleeding disorder Scott Syndrome (12), TMEM16F with its dual functions of ion channel and PL scramblase also mediates membrane wound repair (17), cell signaling and immune responses (18, 19), as well as cell-cell (20, 21) and pathogen-cell fusion (17, 22–24).

Probing the movements of plasma membrane PLs is facilitated by exogenous application of their labeled analogs, which are readily taken up by living cells and incorporated into the plasma membrane with asymmetric distributions resembling those of their endogenous native counterparts (25–32). Fluorescent nitrobenzoxadiazole (NBD)-labeled PLs have been widely used for the determination of the lipid bilayer PL distributions (28). By applying this approach to red blood cells loaded with PLs labeled with NBD or nitroxide spin probe (for electron spin resonance measurements) and extraction of the labeled PLs from the outer leaflet, Williamson and colleagues reported comparable calcium-dependent transmembrane movements for PC, PS, and PE (30). While the approaches involving bovine serum albumin (BSA) back-exchange and the related NBD-fluorescence quenching by BSA have been widely adopted for delineating the endpoint bilayer distribution of PLs, they are not ideal for monitoring PL scrambling because the time frame required for quantitatively extracting BSA-bound NBD-PLs is longer than that of the PL scrambling. In a 1991 landmark paper, McIntyre and Sleight (29) introduced the first fully kinetic scrambling assay using a reducing agent, dithionite, to chemically quench the fluorescence of the outer leaflet residing NBD-PLs. While dithionite quenching has been used widely in proteoliposome systems, its utility in living cells is significantly hampered by interference from plasma membrane transporters such as the Band 3 protein that actively imports dithionite from the extracellular milieu, thereby introducing a quenching signal unrelated to PL scrambling (33). Thus far, all reported NBD-PL-based assays utilize its fluorescence intensity and changes thereof to determine scrambling activities (8, 34). These approaches have collectively cemented the prevailing canon that PL scramblases do not display headgroup selectivity (3, 4, 8, 30, 31, 35–40). Given the limitations of the current repertoire of methods, we devised a new assay involving quantitative assessment of fluorescence polarization (FP) to monitor PL scramblase activity in living cells.

FP is a detection modality that is especially suitable for probing rotational mobility of biomolecules, and has been widely applied for ligand binding and kinase assays (41). In FP experiments (Fig. 1*A*), polarized photons are preferentially absorbed by, and excite, fluorophores whose excitation dipole aligns with the polarization of the photons, leading to a population of excited fluorophores with high orientational coherence. If the fluorophores rotate slowly and emit light before significant randomization of their orientations takes place, the emitted photons retain the polarization of the excitation light beam (vertical as shown in Fig. 1*A*). In contrast, fluorophores with high rotational motility will have their orientations randomized at the time of emission, leading to reduced polarization (depolarization) of the emission. The degree of polarization of the emitted photons can be calculated from the relative intensities (see *Materials and Methods*) measured using two independent and perpendicularly oriented polarization filters. The rotational correlation times of NBD-PLs are comparable to their mean fluorescence lifetimes (both in the nanosecond regime), rendering FP values a sensitive probe for changes in rotational mobility (42–44). Thus, FP measurements provide a clear advantage compared to fluorescence intensity assays in many experimental setups designed to elucidate molecular interactions due to their diagnostic dependence on the rotational mobility, as well as their independence from the fluorescence intensity, which may change for a variety of reasons, including self-quenching (45).

**Figure 1.**
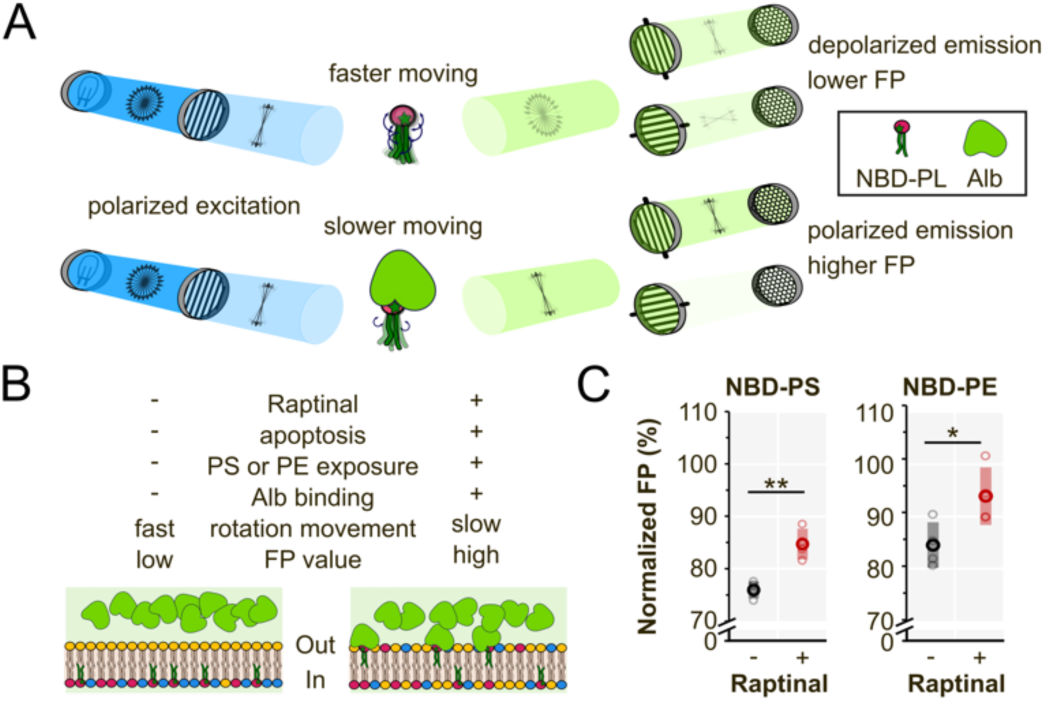
A fluorescence polarization assay for phospholipid movement across bilayers. (A) Simple schematic describing the FP principles. In the initial unpolarized light beam (*blue*), all directions of the individual photon polarizations (*black two-headed arrows*) are represented. An excitation polarization filter that preferentially absorbs photons with horizontal polarization lets through a vertically polarized excitation beam. Excitation photons passing the sample preferentially excite the (possibly mobile) fluorophores that happen to have their excitation dipole vertically oriented, leading to a population of excited fluorophores that have a high orientational coherence. Fluorophores with high rotational mobility (*top*) have their orientations randomized before emission, and the emitted photons (*green*) are highly depolarized. Fluorophores with low rotational mobility (*bottom*), *e.g.,* due to Alb binding, retain their orientations, and the emitted photons retain the high vertical polarization. The emitted photons are passed through independent vertical and horizontal emission polarization filters that preferentially absorb photons with horizontal and vertical polarizations, respectively, and the intensities are measured. (B) Interpretation of FP data in a hypothetical experiment. In non-apoptotic conditions, PS and PE are nearly completely on the inner leaflet of the plasma membrane, inaccessible to Alb on the extracellular side. The rotational mobility of the NBD-PLs is high, and the measured FP value is low. In apoptotic conditions, some NBD-PS and NBD-PE are exposed on the outer leaflet, hallmarks of apoptosis, where they are accessible for Alb binding, and where their rotational mobility is hindered, resulting in a higher net FP value. (C) Normalized Alb-FP values measured immediately after Alb addition from non-apoptotic and apoptotic HeLa cells preloaded with either NBD-PS or NBD-PE. Datapoints from independent replicate experiments (*N* = 3, *light circles*) as well as their means (*dark circles*) ± 1 standard deviation (*shaded box*) are indicated. *P*-values from +Raptinal (apoptotic) vs -Raptinal (non-apoptotic control) comparison (two-tailed Welch’s test): NBD-PS: 0.001, NBD-PE: 0.019. The normalization is based on the first collected datapoint of a kinetic recording. The increased normalized FP signal in apoptotic compared to non-apoptotic cells supports the postulates described in *A* and *B*, indicating that a functional scrambling assay could be designed based on fluorescence polarization. * statistically significant with *p*-value ≤ 0.05, ** statistically significant with *p*-value ≤ 0.01.

Here, we report a cell-based microplate PL scrambling assay whereby FP, rather than fluorescence intensity of NBD-PLs, is monitored, enabling investigations into the headgroup preference of PL scramblases on living cells. By applying this assay on cells with or without endogenous TMEM16F, we deciphered its headgroup preference on the plasma membrane. Coarse-grained molecular dynamics (CGMD) simulations support these findings.

## Results

### A fluorescence polarization assay to distinguish the phospholipid bilayer distribution on the plasma membrane

We hypothesized that the physical binding of albumin (Alb) to NBD-PLs on the outer leaflet of the plasma membrane (46), a prerequisite preceding the extraction and quenching utilized in existing fluorescence intensity-based assays, would retard their rotational mobility and hence, according to well-established physical principles, increase the FP signal (Fig. 1*A*). The degree of increase in the FP signal upon Alb binding would signify the abundance of the NBD on the outer leaflet. If true, this scenario with the fast binding of Alb to outer leaflet residing NBD-PLs, termed Alb-associated FP or Alb-FP, would be an enticing portal for delineating the dynamics of PLs on biological membranes (Fig. 1*B*).

As a litmus test for our hypothesis, we compared the Alb-FP values of NBD-PS and NBD-PE in control and apoptotic HeLa cells, as increases in the extracellularly exposed PS (47) and PE (48) are well-established apoptotic signatures. To elicit the extracellular exposure of PS and PE that are normally located on the inner leaflet of the plasma membrane, we treated cells preloaded with NBD-PS or NBD-PE either with an apoptosis-inducer raptinal (49) or DMSO vehicle for 30 min (the minimum time for raptinal-treated cells to undergo apoptosis). After adding Alb, we immediately detected significantly higher normalized FP values (normalized to a FP reading taken before the inducement of apoptosis) in the apoptotic cells, compared to the non-apoptotic cells (Fig. 1*C*, *red and black datapoints*, respectively). Thus, we confirmed that Alb-FP readings can be used to quantitatively uncouple the NBD-PL signals in the two leaflets.

### Establishing kinetic FP recording for a calcium-stimulated scrambling activity

Encouraged by these results, we proceeded with a fully kinetic FP scrambling assay composed of three stages (Fig. 2*A*). After a baseline FP reading (*blue-shaded region*, typically 1 min), calcium-dependent scrambling was triggered by treating cells with a calcium ionophore, ionomycin, a known pharmacological inducer for TMEM16F scrambling activity (*brown-shaded region*). After a suitable period of plasma membrane scrambling, Alb is added (*green-shaded region*) to selectively reduce the rotational mobility of outer leaflet residing NBD-PLs as well as their ability to subsequently move to the inner leaflet via scrambling or flipping. The FP time-course readings from control and scrambling conditions were normalized to the mean value of their baseline period.

**Figure 2.**
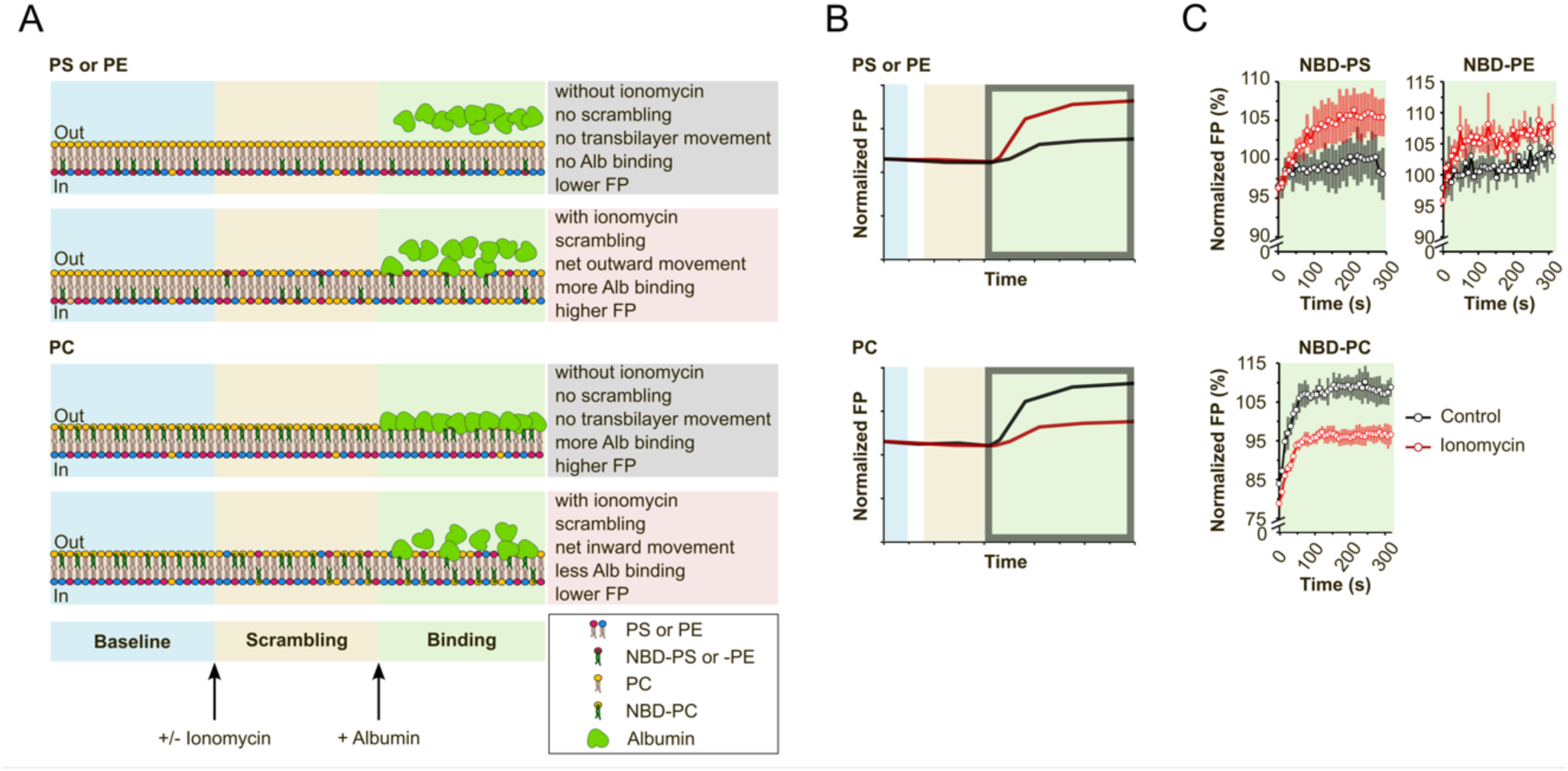
A fully kinetic scrambling assay. (A) Cartoon depiction of a three-stage time-course FP experiment using cells preloaded with NBD-PL: (1) baseline stage (*blue shading*), where baseline FP reading is established, (2) scrambling stage (*yellow shading*), where, depending on the conditions, the asymmetry of PL on the two leaflets is lost, and (3) binding stage (*green shading*), where the binder (Alb) interacts specifically with the extracellularly accessible NBD. In non-scrambling conditions, NBD-PS and NBD-PE are nearly completely located on the inner leaflet of the plasma membrane with high rotational mobility and the measured FP value is low. Adding Alb to the extracellular milieu has little or no effect to the net FP value. In scrambling conditions, there is a net relocation of NBD-PS and NBD-PE to the outer leaflet, where they are accessible to interaction with Alb. The bound NBD-PLs have lower rotational mobility, leading to higher net FP values. In non-scrambling conditions, NBD-PC is mostly located on the outer leaflet, leading to high accessibility for Alb binding and eventually high polarization values. In scrambling conditions, some of the NBD-PC is relocated to the inner leaflet, leading to lower accessibility for Alb binding, and lower net FP values. (B) A cartoon depiction of the expected kinetic FP traces, with black traces indicating non-scrambling conditions and red traces indicating scrambling conditions. A break between the baseline and scrambling periods (*in white*) indicates manual addition of Ionomycin with plate ejected. (C) Alb-FP time-course traces from HeLa cells preloaded with NBD-PS (*N* = 5), NBD-PE (*N* = 5), or NBD-PC (*N* = 5). Means (*circles*) ± 1 standard deviation (*shading*) are indicated. NBD-PS and NBD-PE have lower FP values in non-scrambling (control, *black traces*) conditions compared to the scrambling conditions (*red traces*), indicating net transfer of these NBD-PLs to the outer leaflet. Conversely, NBD-PC has a lower FP value in scrambling conditions compared to the non-scrambling (control) conditions, because in resting conditions there is more NBD-PC on the outer leaflet and upon scrambling, there will be a net transfer to the inner leaflet. The time-course dynamics of the FP have contributions not only from the continuous NBD-PL scrambling and the Alb binding, but also some cellular processes including flipping, the latter of which are also present in the control experiment. After a brief period of increasing FP value, a plateau is reached, indicating saturated binding by Alb. Note that the time axis value 0 refers to the last datapoint in the scrambling stage.

Since FP values alone do not provide the necessary clue to decipher the bilayer distribution of NBD-PL, the interpretation of scrambling events only focuses on the Alb- FP readings (Fig. 2*B*, *green-shaded area*), plotted as a set to infer the direction of the NBD-PL transbilayer movement (Fig. 2*C*).

When we compared the kinetic Alb-FP measurements in the ionomycin-induced scrambling condition and the vehicle-treated control condition in HeLa cells preloaded with NBD-PS, NBD-PE, or NBD-PC, they displayed a change in Alb-FP that matched the predicted models based on their known plasma membrane bilayer distribution (Fig. 2*C*, full time-course traces in *SI Appendix*, Fig. S1). NBD-PS and NBD-PE preloaded cells displayed higher Alb-FP values in scrambling conditions as compared to the non-scrambling control, reflecting a net transfer of NBD-PS and NBD-PE from the inner to the outer leaflet. Conversely, NBD-PC preloaded cells displayed a decrease of Alb-FP values in scrambling conditions as compared to the non-scrambling control, owing to a net transfer of NBD-PC from the outer to the inner leaflet. We confirmed the ability of FP measurements to discern the directionality of PL movement using two additional cell models, U2OS and A549, where ionomycin also triggered a net outward movement of NBD-PS and a net inward movement of NBD-PC (*SI Appendix*, Fig. S2). These data demonstrated that a kinetic FP assay can be used to infer the directionality of NBD-PLs in scrambling conditions.

### The headgroup preference of TMEM16F

After establishing the conditions for the FP scrambling assay, we set forth to address our central question: Does TMEM16F possess headgroup preference *in vivo*? We reasoned that it is possible to deduce whether TMEM16F possesses headgroup preference by examining the scrambling of different PLs in cells with or without endogenous TMEM16F expression. The degree of reduction in the scrambling for a given PL in TMEM16F knock-out (16FKO) compared to WT cells indicates the importance of TMEM16F in the calcium-activated scrambling of that PL. If TMEM16F does not exhibit any headgroup preference for PL scrambling, we would see a universal reduction in transbilayer relocation of all NBD-PLs in 16FKO cells. We started by comparing PS scrambling in NBD-PS preloaded WT and 16FKO HeLa cells. Ionomycin-triggered TMEM16F-dependent PS scrambling is a well-characterized phenomenon detected by annexin-conjugates that are canonical PS-specific probes (12, 13, 50). Unlike the significant increase in Alb-FP values upon ionomycin stimulation of NBD-PS preloaded WT cells, we detected no significant difference in the Alb-FP values of NBD-PS preloaded 16FKO cells with or without ionomycin treatment, indicating minimal NBD-PS outward movement in the absence of TMEM16F expression (Fig. 3*A*, full time-course traces in *SI Appendix*, Fig. S3). Next, we tested the scrambling of NBD-PC and NBD-PE using a similar experimental setup, while adjusting the duration of the scrambling period to account for the differing bilayer distributions and stabilities of these lipids. The Alb-FP values from NBD-PC loaded cells in scrambling and non-scrambling conditions were very different in WT cells compared to 16FKO cells (Fig. 3*B*), indicating a dominant role for TMEM16F in calcium-activated plasma membrane PC scrambling.

**Figure 3.**
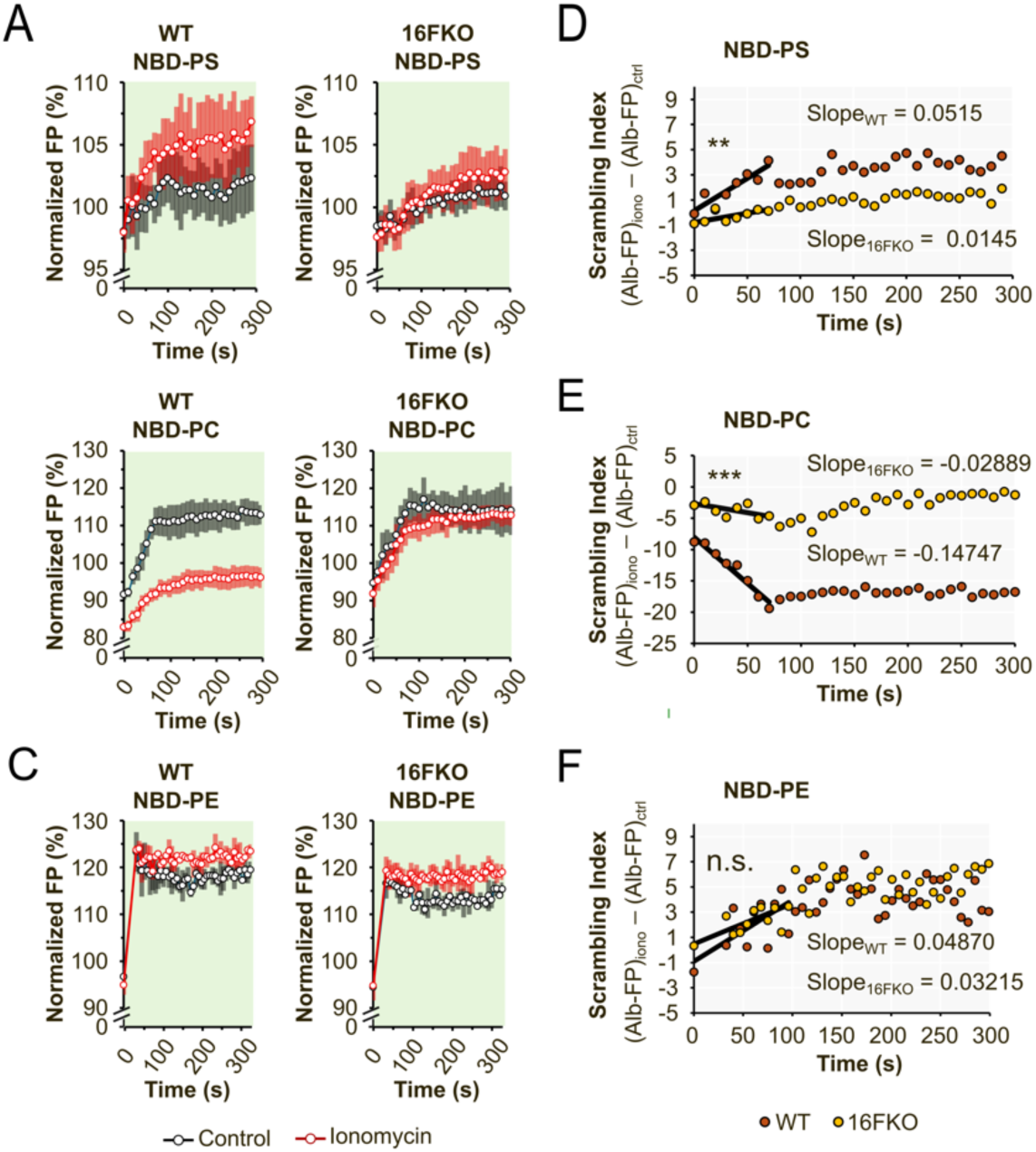
Scrambling index for evaluating TMEM16F scrambling specificity. Alb-FP traces (*left and middle panels*) for WT and TMEM16F HeLa cells were recorded using cells preloaded with (A) NBD-PS (*N* = 7), (B) NBD-PC (*N* = 7), and (C) NBD-PE (*N* = 2) using in both scrambling (*red trace*) and non-scrambling (*black trace*) conditions. Means (*circles*) ± 1 standard deviation (*shading*) are indicated. Scrambling index (*right panels*) is calculated for a given combination of NBD-PL and cellular genotype by subtracting the Alb-FP trace from non-scrambling condition from that of the scrambling condition. The slope calculated from the fitting the first 60 s of the scrambling index trace to a linear function yields the scrambling efficiency for the PL. Mean ± standard error of the slope: (D) NBD-PS, WT: 0.0515 ± 0.0086, NBD-PS, 16FKO: 0.0145 ± 0.0055, 3.54-fold difference in slopes with two-tailed Welch’s test *p*-value 0.0044; (E) NBD-PC, WT: - 0.1475 ± 0.0125, NBD-PC, 16FKO: -0.0289 ± 0.0118, 5.10-fold difference in slopes with *p*-value 1.64e-5; (F) NBD-PE, WT: 0.0487 ± 0.0157, NBD-PE, 16FKO: 0.0321 ± 0.0091, 1.51-fold difference in slopes with *p*-value 0.384. Note that the time axis value 0 refers to the last datapoint of the scrambling stage.

The Alb-FP values from NBD-PE loaded cells in scrambling and non-scrambling conditions in WT cells were similar to those in 16FKO cells (Fig. 3*C*), indicating that removal of TMEM16F does not significantly affect calcium-activated PE scrambling in HeLa cells.

We calculated a “scrambling index” for each PL to obtain a quasi-quantitative view of scrambling effectiveness for comparison between WT and 16FKO cells. The scrambling index for a PL in a given cell type is calculated by subtracting the Alb-FP measurement in non-scrambling conditions from the Alb-FP measurement in scrambling conditions, and fitting the first 60 s, the time frame before significant extraction of NBD-PLs from the outer leaflet of the plasma membrane has taken place (34, 51), to a linear model (see *Materials and Methods*). The ratio in the slopes of the fitted lines indicates the scrambling efficiency for the PL (Fig. 3*D-F*, *SI Appendix*, Fig. S3). For both PS and PC, the scrambling indices of 16FKO cells were significantly different than those of WT cells (Fig. 3*D*, 3.54-fold difference for NBD-PS, *p*-value 0.0044; Fig. 3*E*, 5.10-fold difference for NBD-PC, *p*-value 1.64e-05), indicating a drastic reduction in the calcium-activated PS and PC scrambling at the plasma membranes of cells devoid of TMEM16F expression. In contrast, the scrambling index for PE of WT and 16FKO cells were comparable (Fig. 3*F*, 1.52-fold difference for NBD-PE, *p*-value 0.384), indicating that significant calcium-activated PE scrambling activity at the plasma membrane is retained in the absence of TMEM16F. These data further affirm our findings that TMEM16F displays headgroup preference for PS and PC over PE *in vivo*.

To investigate whether the trends observed in our FP assay hold true at molecular time- and length-scales, we employed coarse-grained molecular dynamics (CGMD) simulations, which have been successfully utilized for evaluating PL scrambling by TMEM16 proteins (52–54). We applied an open conformation structure of calcium-bound TMEM16F (54–56) embedded in a symmetric bilayer with equimolar DOPS, DOPC, and DOPE phospholipids (Fig. 4*A* and 4*B*). By tracking the orientation of individual lipids with respect to the membrane normal in triplicate independent 10 µs simulations (see *Materials and Methods* for details), we detected a distribution of scrambling events that are in agreement with those from our FP assays, whereby TMEM16F exhibits a significant preference for PS and PC over PE (Fig. 4*C*).

**Figure 4.**
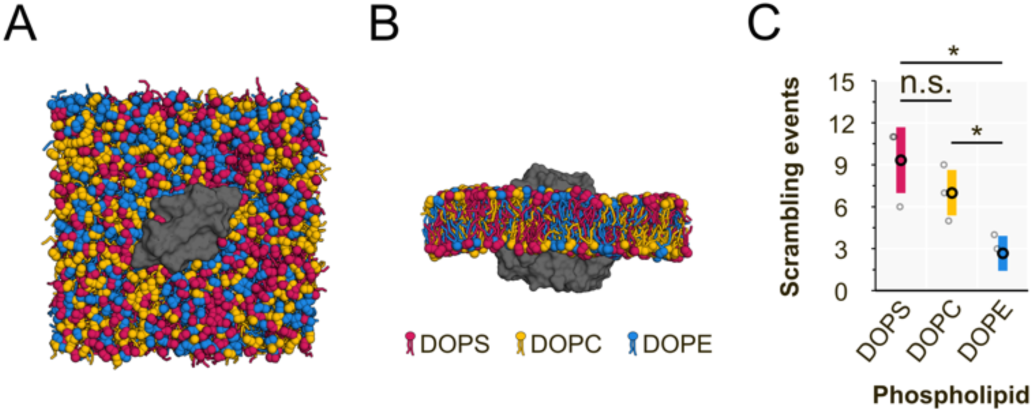
Coarse-grained molecular dynamics simulation of TMEM16F embedded in an equimolar and symmetric bilayer of DOPS, DOPC, and DOPE phospholipids. (A) Top, and (B) side view of the initial frame of the simulation. (C) The total number of the scrambling events for DOPS, DOPC, and DOPE in the last 9 μs of CGMD simnulation. Datapoints from independent replicate experiments (*N* = 3, *light circles*) as well as their means (*dark circles*) ± 1 standard deviation (*shaded box*) are indicated. *P*-values from two-tailed Welch’s tests: DOPS *vs*. DOPC: 0.321 (n.s.), DOPS *vs*. DOPE: 0.038 (*), DOPC *vs*. DOPE: 0.044 (*). n.s., not significant with p-value > 0.05; *, statistically significant with *p*-value ≤ 0.05.

In summary, our cell-based FP scrambling assays, together with CGMD simulations, provide evidence for PL headgroup preference of TMEM16F. Taken together, these findings support the notion that this calcium-activated PL scramblase preferentially scrambles PC and PS over PE.

## Discussion

The question concerning headgroup preference of PL scramblases is relevant to the burgeoning field of lipid scrambling and membrane biology in general. While there is a consensus in the published literature that phospholipid scramblases do not exhibit headgroup preference (3, 4, 8, 30, 31, 35–40), we wanted to critically revisit this question given the limitations in currently available detection methods. Thus, we designed an approach based on fluorescence polarization that is applicable to living cells. In contrast to the prevailing views, we found that TMEM16F displays headgroup preference for PS and PC over PE, a discovery supported by CGMD simulations.

Our scrambling assay employs the same tail-derivatized NBD-PLs that have been utilized in prior PL scrambling assays, which rely on the selective elimination of the NBD-PL fluorescence located on the outer layer of the plasma membrane, by either dithionite reduction of NBD to its non-fluorescent derivative (29), or physical removal of NBD-PL by BSA (34). The dithionite quenching has the advantage of enabling real-time recording, yet it suffers from a high degree of false-positive signal from dithionite internalization, where it can act on the inner leaflet-residing NBP-PLs as well (29, 57).

While the BSA back-exchange (also called quenching) is specific for the NBD-PLs on the extracellular leaflet, the kinetics of NBD-PL removal are much slower than the scrambling and is often coupled with a physical isolation step for accurate measurements at a single time point (34), rendering this approach more suitable for probing steady-state PL bilayer distributions.

We opted for detection of fluorescence polarization (FP) to capture the rotational mobility of the NBD-PLs rather than probing for their fluorescence intensity, and we applied Alb to the extracellular media to specifically reduce the mobility of the outer leaflet-residing NBD-PL. Hence, with FP measurement, we attain temporal resolution comparable to the dithionite-based method, while also achieving the spatial specificity that can be afforded by BSA back-exchange/quenching. Using a kinetic recording mode on a plate reader, we could capture the change in Alb-FP signals correlated with directional movements of NBD-PLs as a readout of PL scrambling. We adjusted the durations of the scrambling period in our measurements for different PLs to accommodate their different resting bilayer distributions and stabilities on the plasma membrane. We allowed ionomycin to act on NBD-PS loaded cells for 2 min before Alb addition. Given the tendency of NBD-PE for metabolization (58) (that may be triggered by ionomycin stimulation (59)) and internalization to organelle membranes, we shortened the NBD-PE scrambling duration to 30 sec. Since PC predominantly resides on the outer leaflet in resting cells, we extended the scrambling duration to ∼4-9 min for a significant amount of PC to achieve inward movement before adding Alb. In view of the intrinsic physical complexities behind the FP signal, especially in a cell-based assay, our assay is currently not fully quantitative. Nonetheless, comparing the scrambling indices between WT and 16FKO cells enabled us to decipher the ionomycin-mediated bilayer movement of PS, PE, and PC and extrapolate the headgroup preference of TMEM16F. In addition to confirming that TMEM16F is the dominant calcium-activated PS and PC scramblase on the plasma membrane that responds to ionomycin treatment, we detected significant retention of NBD-PE scrambling activity in 16FKO cells, indicating TMEM16F is not the major PE scramblase in HeLa cells.

Although we performed parallel control experiments in our study, we acknowledge the undefined composition and complex PL dynamics in living cells utilized in our experiments. We also need to keep in mind that several other known processes besides scrambling, including ATP-dependent restoration of the plasma membrane asymmetry by flippases and floppases, PL trafficking and catabolism, as well as the Alb binding to the NBD-PLs, contribute to the kinetics of the measured FP values.

Importantly, our findings on the headgroup preference of TMEM16F on the plasma membrane are supported by CGMD simulations of TMEM16F in a bilayer with equal proportions of DOPS, DOPC, and DOPE, thereby showing that the preference of TMEM16F for PS and PC over PE extends to molecular length- and timescales: we observed ∼4- and 3-fold preferences in our cell-based FP scrambling assay (Fig. 3*D*-*F*) and in CGMD simulations (Fig. 4*C*), respectively.

While our *in vivo* experimental measurements and simulation results demonstrate the headgroup preference of TMEM16F, we need to consider the reasons that may contribute to the difference between our work and the previously reported *in vitro* study (38). Watanabe and colleagues reconstituted purified murine Tmem16f onto a synthetic lipid bilayer array and measured the recovery of fluorescence signals after photobleaching TopFluor-TMR-conjugated PLs that are spiked into the bilayer (38).

Using this elegantly designed platform, they recorded comparable scrambling rates when TopFluor-TMR-conjugated PS, PE or PC were introduced into the bilayer. One major difference from our studies is the relatively simple and skewed chemical composition of PLs on the reconstituted lipid array, where the spiked-in analytes, TopFluor-TMR-conjugated PLs, are surrounded by a disproportionate amount of POPC. Given the tendency of PLs to undergo self-sorting according to their headgroup chemical and physical properties (*e.g.*, the formation of lipid microdomains on the cellular plasma membrane (60)), the scrambling capacity of Tmem16f on the reconstituted lipid array may not reflect its intrinsic nature under physiological conditions. While the lipid composition in our CGMD is also artificial, the presence of three major PLs in equal amounts imposes a spatial competition among PL headgroups for interaction and scrambling by TMEM16F that is absent in the reconstituted lipid array experiment reported by Watanabe and colleagues.

With the discovery of the scrambling preference of TMEM16F for PC and PS on the plasma membrane, we believe this study will open many avenues for future research that could be best carried out using FP measurements either on native membranes of living cells or in reconstituted systems. These questions include: What are the structural determinants conferring PL headgroup preference to TMEM16F? Which scramblase(s) preferentially act on PE and possibly other PLs? Do other scramblase families, such as the Xkr, TMC and TMEM63 families, also display PL headgroup preference? Which proteins, lipids, and small molecules modulate membrane scrambling in physiological conditions? We anticipate that this FP scrambling assay can complement the existing assays on proteoliposomes and other reconstituted systems that provide useful model systems. This new experimental approach may also pave the way for uncovering the underlying mechanisms of organellar membrane scrambling.

## Materials and Methods

### Reagents

We purchased 18:1-06:0 phospholipids NBD-PC (cat# 810132), NBD-PS (cat# 819194), and NBD-PE (cat# 810155) from Avanti Polar Lipids, dissolved them in anhydrous DMSO (Invitrogen, D12345) to 0.2 mM (100× stock), and stored at -20 °C as single-use aliquots. We purchased ionomycin from Cayman Chemical (cat# 10004974) supplied at 14.1 mM in ethanol, diluted it to 5 mM with 100% ethanol, and stored it at -20 °C as single-use aliquots. We purchased Raptinal (cat# 9626-10) from BioVision, prepared it in anhydrous DMSO to 5 mM (500× stock), and stored at -20 °C as single-use aliquots. Gelatin solution (cat# ES-006-B) used for cell culture plate coating was purchased from EMD Millipore Sigma. We used fatty acid-free BSA from Gemini Bio-Product (cat# 700- 107p) in the initial experiments and later switched to recombinant human albumin (10% w/v ALBIX) from Albumedix.

### Cell culture

We purchased STR profile-certified HeLa and U2OS cells from the UCSF Cell and Genome Engineering Core facility. A549 cells were a gift from Dr. Yu-Ting Chou (Trever Bivona laboratory, UCSF). Cells were cultured in 37 °C incubators supplied with 5% CO_2_. Clonal TMEM16F CRISPR-Cas9 knock-out HeLa cells were generated, screened, and validated as described in detail for another human cell line (61). We cultured the HeLa and U2OS cells in DMEM media (Gibco, 10569044) supplemented with 10% FBS (Cultra Pure US origin, Axenia BioLogix), and A549 cells in advanced DMEM media (Gibco, 12491-015) supplemented with 2% fetal bovine serum (Gibco, 26140-079), 1× GlutaMAX (Gibco, 35050-061), and 100 μg/ml Normocin (InvivoGen, ant-nr-05). Cells were dislodged for passaging with TrypLE (Gibco, A1217701) after washing twice with ambient temperature DPBS (Gibco, 14190136). We froze these cells in Recovery Cell Culture Freezing Medium (Gibco, 12648010) in liquid nitrogen. We used MycoStrip (InvivoGen, rep-mys) to test for mycoplasma contamination and Cell Culture Contamination Detection Kit (Invitrogen, C7028) to test for fungi and mold contamination.

### Fluorescence polarization scrambling assay

We seeded cells on gelatin-coated 96-well flat-bottom black plates (Greiner, 655090) at a sufficient cell density to produce 80 to 90% confluency for the next day’s experiment. Due to size and growth rate differences, the following seeding densities (10^4^ cells per well) were used: HeLa: 0.8, U2OS: 1.2, A549: 1.6. Based on our experience, cells that are either too dense or too sparse would affect the FP reading and thus compromise experimental accuracy. Additionally, viable, healthy and contaminant-free cells (mycoplasma and fungi/mold) were found to be essential for these experiments.

Experiments were carried out with fluorescence polarization-equipped BMG Labtech PHERAstar *FSX* and Agilent Synergy H4 microplate readers. Considering the data collection rate of these instruments in kinetic FP mode, we only plated cells to 12 (PHERAstar *FSX* experiments) or 4 (Synergy H4 experiments) adjacent wells of a 96- well plate for a single experiment so that the FP was recorded for every 7 to 10 s for a given well. In each experiment, the plate layout was as follows: cells in all but two blank wells received NBD-PL. Both blank and NBD-PL wells were split into two groups with an equal number of wells for apoptosis and non-apoptosis or scrambling and non-scrambling control conditions. First, the cells were washed three times (150 μL each) with Buffer W (HBSS (Gibco, 14065056) supplemented with 10 mM HEPES (Fisher Scientific, BP299-100)) and incubated with 100 μL of 0.2 μM NBD-PL in Leibovitz’s L-15 medium (Gibco, 1141504) for 20 min at ambient temperature covered from light. After two washes (150 μL each) with Buffer W, we added 100 μL of Buffer H (1x HBSS, 20 mM HEPES, 20 mM glucose (Fisher Scientific, BP350-1), 1 mM sodium pyruvate (Sigma-Aldrich, P2256), 1.8 mM CaCl_2_, (Fisher Scientific, C79-500)) to each well. For apoptosis experiments, 10 μM raptinal or DMSO are included in the Buffer H. We then placed the plate in the plate reader and launched the preset method with parameters listed below. For scrambling experiments, we manually added 5 μL of 42 μM ionomycin (scrambling conditions) or ethanol vehicle (non-scrambling conditions) diluted in Buffer H during the pause cycle/step as specified below. Ionomycin and control working solutions were prepared freshly for each plate. We employed the built-in injector to add 10 μg ALBIX to each well in experiments performed on the PHERAstar *FSX*, or manually pipetted 10 μg of BSA to each well in experiments performed on the Synergy H4. The albumins were prepared in buffer H and filtered through a 0.45 μm surfactant-free cellulose acetate Nalgene syringe filter prior to use (723-254{Citation}5 ThermoFisher Scientific). We found no significant difference in data quality between recombinant albumin and BSA used in the assays. We chose recombinant albumin in the later experiments. Data acquisition parameters are listed below.

PHERAstar *FSX* from BMG Labtech operated with the PHERAstar software (v5.70 R5) was used to collect data for Fig. 1*C*, 2*C*, 3*A*, 3*B*, and *SI Appendix*, Fig. S1 to S3 with the following details. Optical settings: FP module: 488_Ex_, 520_EmA_ 520_EmB_, kinetic plate mode, focal adjustment enabled, gain adjustment mP = 180. General settings: bottom optic, settling time 0.1 s, target temperature 37 °C, number of flashes = 100.

Injection settings: 40 μL volume (equivalent to 10 μg of ALBIX), 100 μL/s injection speed, standard injection spoon (type A1), smart dispensing option enabled. Shake settings: after the pause cycle #, at 100 rpm for 2 s with the double orbital option.

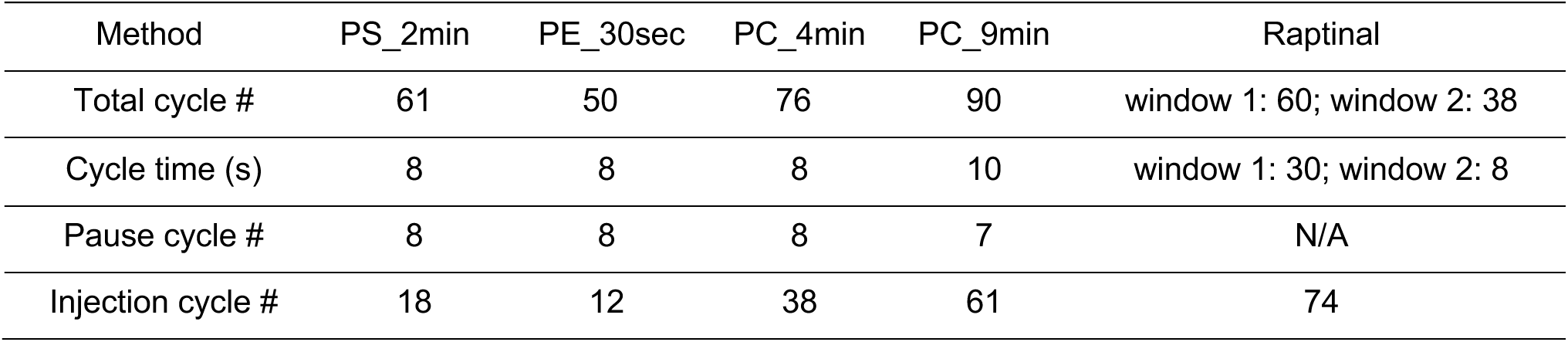

| Method | PS_2min | PE_30sec | PC_4min | PC_9min | Raptinal |
| --- | --- | --- | --- | --- | --- |
| Total cycle # | 61 | 50 | 76 | 90 | window 1: 60; window 2: 38 |
| Cycle time (s) | 8 | 8 | 8 | 10 | window 1: 30; window 2: 8 |
| Pause cycle # | 8 | 8 | 8 | 7 | N/A |
| Injection cycle # | 18 | 12 | 38 | 61 | 74 |

Synergy H4 from Agilent with Gen5 (v2.09) was used to collect data in Fig. 3*C* with the following settings. Detection method: Fluorescence polarization. Excitation: 485/20 nm; Emission: 528/20 nm. Optic position: Top, 510 nm, Gain: 80, Read speed: Normal, Read height: 4 mm. Light source: Xenon flash. Read type: Kinetic plate mode with three windows. Kinetic window 1: 1 min (7 s interval, 9 reads); plate out (manually added 5 μL of 42 μM ionomycin); shake for 2 s at medium speed; kinetic window 2: 30 s (7 s interval, 5 reads); plate out (manually added 100 μL of 100 μg BSA); kinetic window 3: 5 min (7 s interval, 43 reads).

Fluorescent background signals from the wells without NBD-PL scrambling and non-scrambling controls were taken into account via preselecting them as Blanks in the method acquisition file. For data acquired on the PHERAstar *FSX*, the calculated mP values from the PHERAstar analysis MARS software (version 4.01 R2) were exported to the Apache OpenOffice Spreadsheet.

For data acquired on the Synergy H4, raw reads of fluorescent signals were exported to Microsoft Excel, and FP values were manually calculated using the following equation:

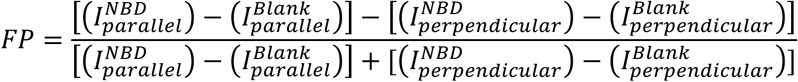

Where *I* is fluorescence intensity, superscripts *NBD* and *Blank* refer to the cell preloading, and subscripts *parallel* and *perpendicular* refer to the emission filter orientation.

The FP values were normalized to the mean of the baseline stage before statistical analyses and plotting.

### Data fitting for Scrambling Index

Scrambling index (SI) was calculated by subtracting FP-Alb trace from non-scrambling condition from that of the scrambling condition for each experiment. The first 60 s of the resultant trace are fitted to a line using lm function in R, and the slope was extracted as a measure of scrambling efficiency for that PL. The slope ratio was calculated between the scrambling and non-scrambling conditions, as were the statistical significance using two-sided Welch’s test in R.

### Coarse-grained molecular dynamics simulation

A symmetric model of the TMEM16F dimer with dilated TM4-TM6 grooves in both subunits was created by copying and superposing the chain with an “open groove” conformation that resulted from extensive all-atom simulations (cluster 10 in ref (56)). Coarse-grained systems were prepared as described previously (54). Briefly, *martinize2* (62) and *insane* (63) scripts were employed to embed the CG protein into a symmetric and equimolar DOPC:DOPS:DOPE bilayer. This process was repeated independently for three replicates to ensure randomized placement of the PL molecules in each system.

CGMD simulations were performed using the Marini 3.0.0 force field (64) and Gromacs 2023.3 (65) using a 20 fs time step. Non-bonded interactions were described by reaction-field electrostatics and Van der Waals potentials, using a 11 Å cut-off. Constant temperature (T = 310 K, τ_T_ = 1 ps) and pressure (P = 1 bar, τ_T_ = 12 ps) were maintained using the velocity rescaling thermostat (66), and semi-isotropic Parrinello-Rahman barostat (67), respectively. After energy minimization and a short NPT equilibration, all systems were simulated for 10 μs. The first 1 μs was considered a system equilibration phase and omitted from analyses.

PL scrambling was analyzed from CGMD simulations as described previously (53, 54, 68). Briefly, for every PL molecule in every simulation frame, the angle ϑ between the director vector and the membrane normal was measured. After smoothing the data using a 100 ns running average, a scrambling event is recorded when ϑ transitions from the outer leaflet value (between 145° and 180°) to the inner leaflet value (between 0° and 35°) or *vice versa*.

## Data, Materials, and Software Availability

The raw data presented and R code used in this manuscript are available from the authors upon request.

## Acknowledgements

We thank Dr. Edward J. (EJ) Dell from the BMG Labtech for helpful discussions and for customizing the built-in injectors on PHERAstar *FSX* to suit our assays, Dr. John M. Gilchrist for sharing his extensive knowledge on dithionite quenching assay, Dr. Yu-Ting Chou for gifting the A549 cells, Dr. Ya-Chu Chang for critical reading of this manuscript, Prof. Dean Sheppard and his lab members, especially Amha Atakilit and Xin Ren, for generously sharing cell culture resources, Marena Tynan-La Fontaine, Armando Martinez, Hongbin (Ben) Yuan and Xingnu (Jessie) Zhai for their continuing logistics support in the Jan lab. Y.N.J and L.Y.J are investigators at the Howard Hughes Medical Institute. This work is supported by NIH grants R35NS122110 to LYJ, R35NS137312 to YNJ, and R01GM137109 to MG. DC is supported by Le Studium Fellowship Loire Valley Institute for Advanced Studies (Tours & Orléans, France), Cancéropôle Grand-Ouest IMACALSEQCAP and Ligue Contre le Cancer “Grand-Ouest”.

## Author Contributions

C.F.T. and S.T.T. conceived the project. C.F.T. designed and executed all experiments. S.T.T. independently validated the experiments. N.v.H. performed simulations. D.C. performed data fitting. C.F.T., S.T.T., N.v.H., and D.C. interpreted the data. C.F.T., S.T.T., and L.Y.J. wrote the manuscript. All authors edited the manuscript. M.G., Y.N.J., and L.Y.J. oversaw and supervised the project and provided the project resources.

## Supporting Information

**Fig. S1.**
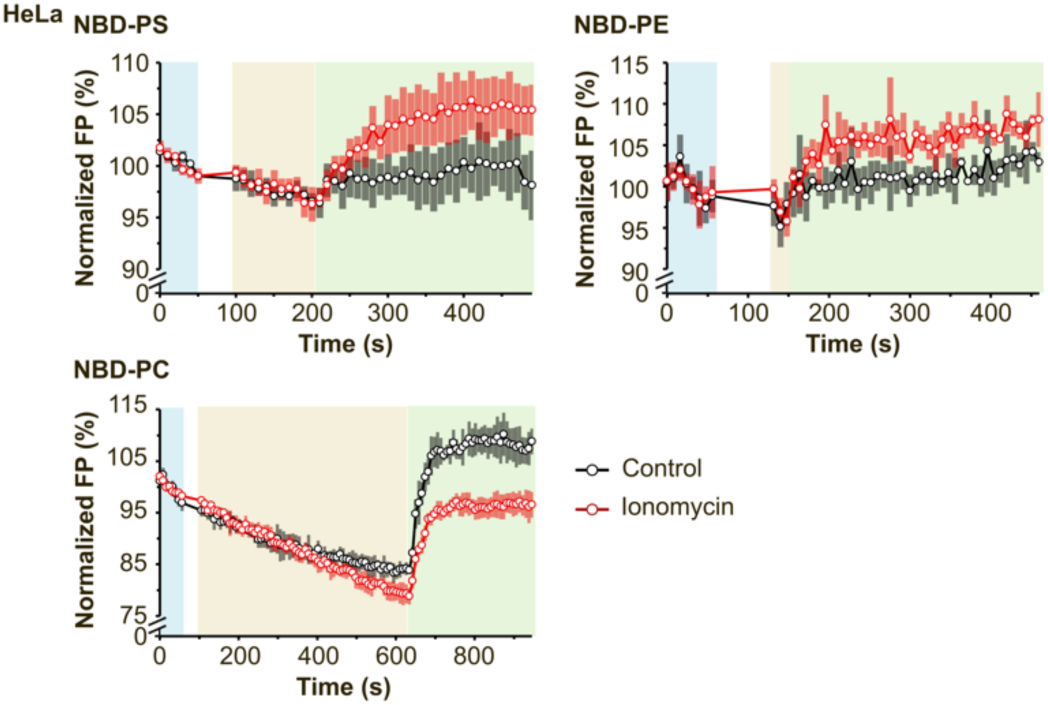
Full time-course data of the kinetic calcium-induced phospholipid scrambling in WT HeLA cells. The cells were preloaded with either PC, PS, or PE, and treated by either ionomycin to induce scrambling (*red traces*) or left as controls with vehicle treatment (*black traces*). For each trace, *N* = 5. Means (*circles*) ± 1 standard deviation (*shading*) are indicated.

**Fig. S2.**
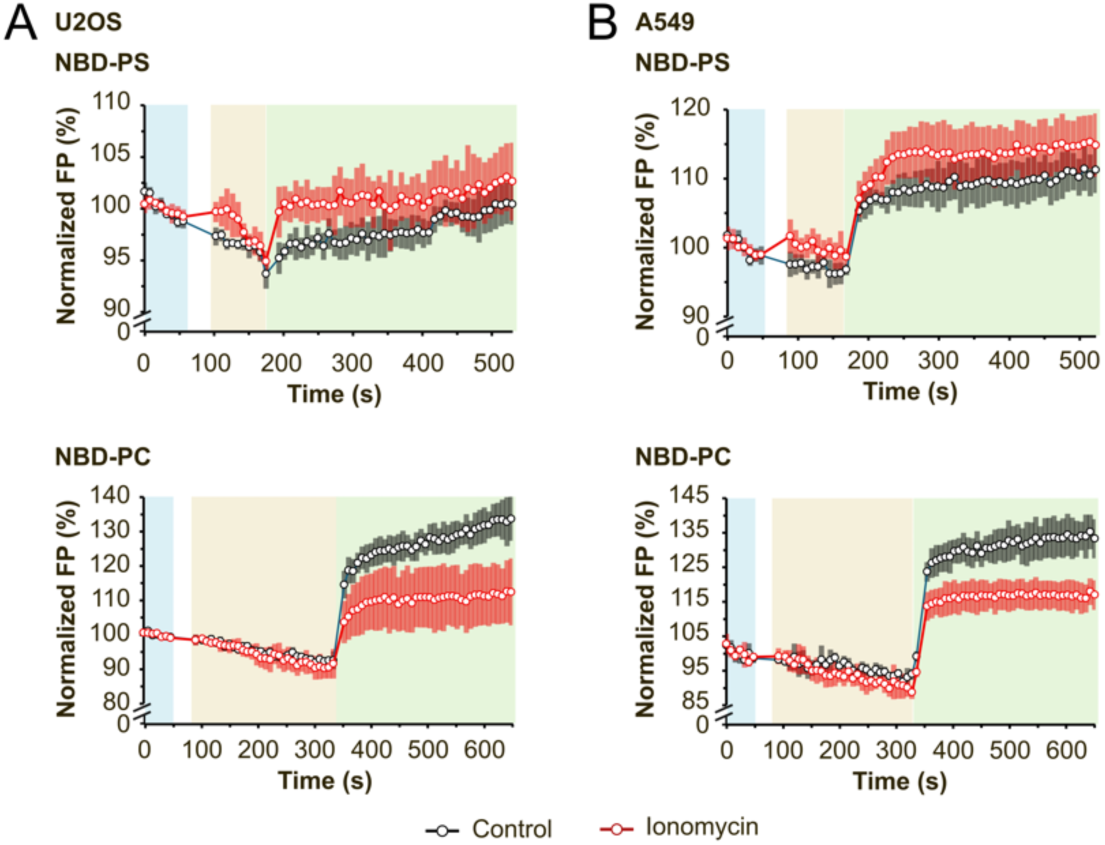
The ability of the FP assay to discern the directionality of PL movement on the plasma membrane was verified using two additional cell lines, (A) U2OS and (B) A549. The data from both cell lines corroborates the findings in HeLa cells: In NBD-PS preloaded cells, the FP signal from scrambling conditions (*red traces*) is higher than in the control (non-scrambling, *black traces*) conditions, and in NBD-PC preloaded cells, the FP signal from scrambling conditions (*red traces*) is lower than in the control (non-scrambling, *black traces*) conditions. For each trace, *N* = 5. Means (*circles*) ± 1 standard deviation (*shading*) are indicated.

**Fig. S3.**
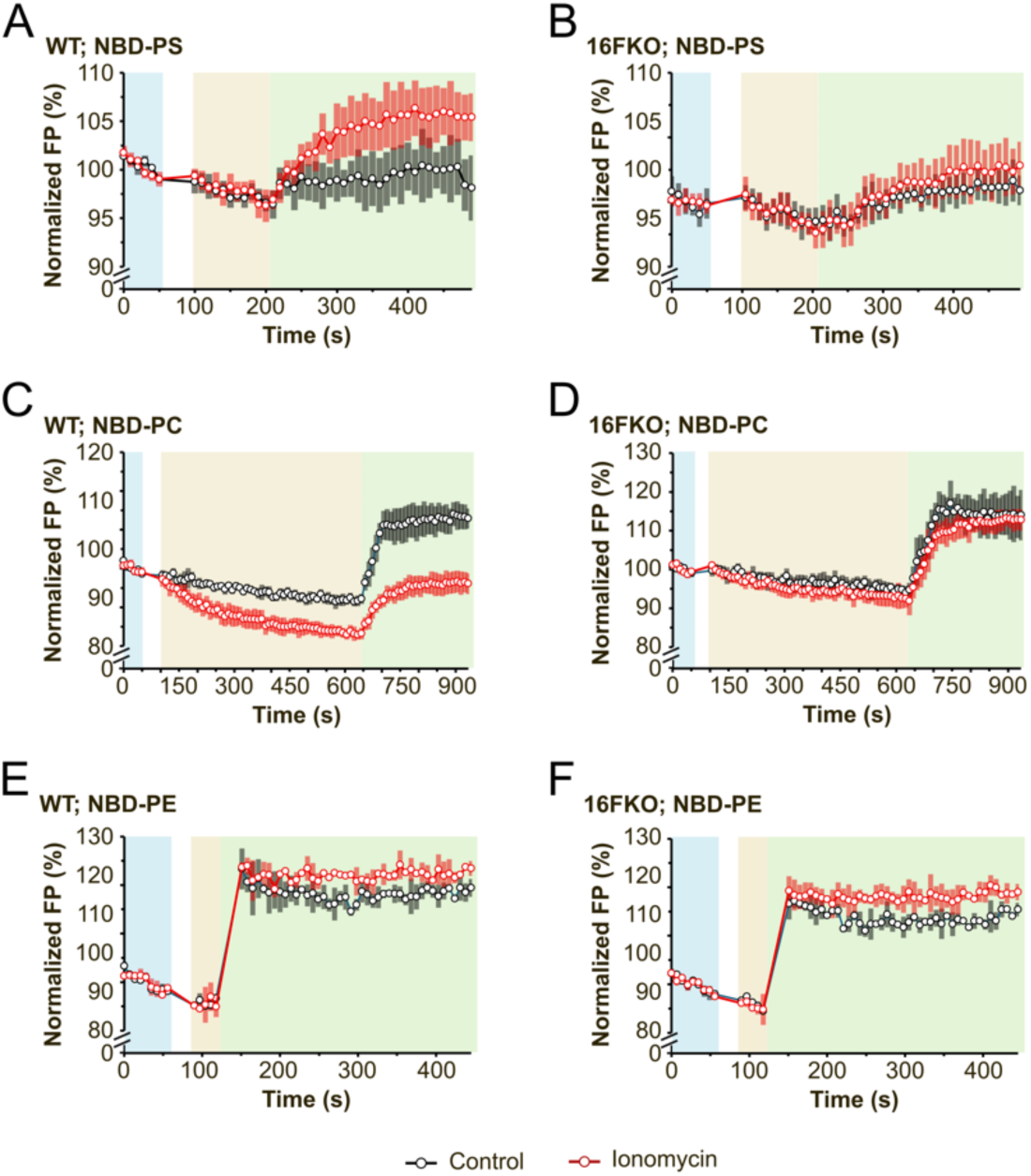
Full time-course data of the kinetic calcium-induced phospholipid scrambling in WT (A, C, E) and 16FKO (B, D, F) HeLa cells. In all cases, the difference between scrambling (red *traces*) and non-scrambling (*black traces*) conditions is narrower in the 16FKO cells compared to the WT cells. For A and B (NBD-PS), *N* = 7, for C and D (NBD-PC), *N* = 7, and for E and F (NBD-PE), *N* = 2. Means (*circles*) ± 1 standard deviation (*shading*) are indicated. In NBD-PC and NBD-PS preloaded cells, the differences are nearly abolished, indicating that TMEM16F contributes to most of the calcium-activated PC- and PS-scrambling activities in HeLa cells. The insignificant difference between scrambling (red *traces*) and non-scrambling (*black traces*) conditions in NBD-PE preloaded 16FKO cells and WT cells suggests a relatively minor effect on calcium-activated PE scrambling activity in cells lacking TMEM16F.

